# Hundreds of nuclear and plastid loci yield insights into orchid relationships

**DOI:** 10.1101/2020.11.17.386508

**Authors:** Oscar Alejandro Pérez-Escobar, Steven Dodsworth, Diego Bogarín, Sidonie Bellot, Juan A. Balbuena, Rowan Schley, Izai Kikuchi, Sarah K. Morris, Niroshini Epitawalage, Robyn Cowan, Olivier Maurin, Alexandre Zuntini, Tatiana Arias, Alejandra Serna, Barbara Gravendeel, Maria Fernanda Torres, Katharina Nargar, Guillaume Chomicki, Mark W. Chase, Ilia J. Leitch, Félix Forest, William J. Baker

**Author notes:** Joint senior authors (Mark W. Chase, Ilia J. Leitch, Félix Forest, William J. Baker).

## Abstract

**Premise of the study:** Evolutionary relationships in the species-rich Orchidaceae have historically relied on organellar DNA sequences and limited taxon sampling. Previous studies provided a robust plastid-maternal phylogenetic framework, from which multiple hypotheses on the drivers of orchid diversification have been derived. However, the extent to which the maternal evolutionary history of orchids is congruent with that of the nuclear genome has remained uninvestigated.

**Methods:** We inferred phylogenetic relationships from 294 low-copy nuclear genes sequenced/obtained using the Angiosperms353 universal probe set from 75 species representing 69 genera, 16 tribes and 24 subtribes. To test for topological incongruence between nuclear and plastid genomes, we constructed a tree from 78 plastid genes, representing 117 genera, 18 tribes and 28 subtribes and compared them using a co-phylogenetic approach. The phylogenetic informativeness and support of the Angiosperms353 loci were compared with those of the 78 plastid genes.

**Key Results:** Phylogenetic inferences of nuclear datasets produced highly congruent and robustly supported orchid relationships. Comparisons of nuclear gene trees and plastid gene trees using the latest co-phylogenetic tools revealed strongly supported phylogenetic incongruence in both shallow and deep time. Phylogenetic informativeness analyses showed that the Angiosperms353 genes were in general more informative than most plastid genes.

**Conclusions:** Our study provides the first robust nuclear phylogenomic framework for Orchidaceae plus an assessment of intragenomic nuclear discordance, plastid-nuclear tree incongruence, and phylogenetic informativeness across the family. Our results also demonstrate what has long been known but rarely documented: nuclear and plastid phylogenetic trees are not fully congruent and therefore should not be considered interchangeable.

## INTRODUCTION

Phylogenetic inference based on nuclear and organellar DNA sequences has revolutionised plant systematics and evolution (Cameron et al., 1999; Soltis et al., 2000; Eiserhardt et al., 2018). From species complexes (Bogarín et al., 2018; Fernández-Mazuecos et al., 2018; Pérez-Escobar et al., 2020) to families and beyond (Bateman et al., 2018; Nauheimer et al., 2018; Wan et al., 2018; Wong et al., 2020), molecular phylogenetics has radically shaped our understanding of plant evolution at widely varying scales and subsequently had a drastic effect on their classification in order to maintain monophyletic groups (Chase et al., 2016). Historically, Sanger-sequenced DNA markers have been widely used to infer phylogenies at various taxonomic levels (Baldwin, 1992; Baldwin and Markos, 1998; Cameron et al., 1999; Soltis et al., 2000). In particular, plastid DNA loci such as *rbcL, matK*, and the *trnL-trnF* region, the nuclear ribosomal internal transcribed spacer (nrITS) and the low-copy nuclear gene *Xdh* have been frequently used/studied (Chase et al., 1993; Baldwin et al., 1995; Soltis et al., 1999; Hilu et al., 2003; Górniak et al., 2010). This is because of the relative ease of sequencing these DNA regions, particularly those in high-copy numbers (plastid regions and nrITS, but see Górniak et al. 2010).

Recently, implementation of high-throughput DNA sequencing methods has expanded the number of DNA regions available for phylogenetic inference, but as with the earlier studies there has been a continuing focus on plastid genes (Edger et al., 2018; Li, Li, et al., 2019) and plastomes (Ross et al., 2016; Guo et al., 2017; Li, Yi, et al., 2019). The rapid growth of organellar phylogenomic studies is an evident trend in many plant families including Araceae (Henriquez et al., 2014), Berberidaceae: (Sun et al., 2018), Lauraceae (Song et al., 2020), Leguminosae (Zhang et al., 2020) and Orchidaceae (Givnish et al., 2015) due to the relative ease with which they are generated: parallel sequencing at shallow coverage (less than 1X) of genomic library preparations derived from recent or historical plant material can yield millions of organellar DNA sequencing reads for hundreds of individuals. This enables the sequencing of dozens of plastid loci at a lower cost per sample than when using Sanger sequencing (Straub et al., 2012; Dodsworth, 2015).

With at least 25,000 species and 700 genera, Orchidaceae are one of two largest angiosperm families, distributed across most terrestrial biomes. Orchids display a wide range of vegetative and reproductive traits that have long captivated biologists. They exhibit unusual relationships with animal pollinators (Darwin, 1877; Jersáková and Malinová, 2004; Ramirez et al., 2011; Martins et al., 2018) and mycorrhizal fungi (Dearnaley, 2007; Rasmussen, 2015; Fochi et al., 2017), which have been the subject of detailed studies. Other unusual characters of interest include the velamen, a tissue fostering water uptake and protecting orchid roots in epiphytic orchids (Zotz and Winkler, 2013; Chomicki et al., 2015) and their mostly anemochorous (Arditti and Ghani, 2000; Barthlott et al., 2014) or sometimes animal-dispersed seeds (ants, bats, bees, crickets and frugivorous birds; (Suetsugu et al., 2015; Morales-Linares et al., 2018). Their global distribution, high species-richness in the tropics and wide variety of functional traits and ecological interactions makes them an excellent model group for studying how biotic and abiotic factors affect their diversification (Givnish et al., 2015, 2016; Pérez-Escobar, Chomicki, Condamine, de Vos, et al., 2017; Pérez-Escobar, Chomicki, Condamine, Karremans, et al., 2017).

Understanding orchid relationships is essential to enable the interpretation of their extraordinary diversity, and as a result orchid phylogenetics have been investigated intensively (Dressler, 1993; Cameron et al., 1999; Freudenstein and Chase, 2015; Givnish et al., 2015). By including representatives of nearly all major taxa, phylogenetic trees inferred from the analysis of mostly organellar loci have provided a robust set of relationships for most orchid tribes and subtribes (reviewed in (Chase et al., 2015)). These well-supported organellar phylogenetic frameworks have been subsequently employed to infer relationships at different taxonomic levels and investigate the historical biogeography and evolution of selected traits for over 20 years (Neyland and Urbatsch, 1996; Cameron et al., 1999). However, the extent to which the evolutionary history of the uni-parentally inherited organellar genomes (Chang et al., 2000; Cafasso et al., 2005) track that of the bi-parentally inherited and recombinant nuclear genome in the orchid family has not been properly tested in a phylogenomic framework. In fact, there is mounting evidence of incongruence driven by biological phenomena including hybridization and incomplete lineage sorting across the angiosperms (Soltis and Kuzoff, 1995; Smith et al., 2015; Pérez-Escobar et al., 2016; Vargas et al., 2017; Schley et al., 2020), and the extent to which it prevails in orchids remains to be assessed.

In this study, we generate a phylogenomic nuclear dataset for Orchidaceae relying on the universal Angiosperms353 target capture probe set (Johnson et al., 2019). For this, we sampled 75 species representing all the five subfamilies and the majority of the tribes. To assess nuclear-plastid intra- and intergenomic topological conflict, we also present a plastid phylogenomic tree inferred from the published sequences of 78 genes for 264 species, also representing all orchid subfamilies, and most of the recognised tribes (Chase et al., 2015). Lastly, we compared the phylogenetic informativeness of nuclear and plastid loci. We address the following topics: i) the extent to which the maternally inherited orchid plastid tree is congruent with that of the bi-parentally inherited nuclear genome and ii) how well these low-copy nuclear genes perform in recovering strongly supported relationships compared to plastid genes/genomes.

## MATERIALS AND METHODS

### Taxon sampling

We included 75 orchid species representing 69 genera, 24 subtribes (out of 46 currently accepted; Chase et al., 2015), 16 tribes (out of 21), and all five subfamilies. In addition, fourteen species from non-orchid monocot families were included as outgroup taxa (**Appendix S1**). Newly generated nuclear DNA data were produced from 62 vouchered accessions stored in the DNA and tissue bank of the Royal Botanic Gardens, Kew (https://dnabank.science.kew.org/homepage.html). These DNA samples had been previously extracted from silica-dried leaves using a modified CTAB method (ref). Each sample has a voucher herbarium specimen hosted in the Kew herbarium (K; **Appendix S1**). To expand our taxon sampling, nuclear short Illumina sequencing reads were datamined from the Sequence Read Archive (SRA) using the fastq-dump software of the SRAtool-kit package (available at https://ncbi.github.io/sra-tools/install_config.html), and from the 1KP data repository (https://sites.google.com/a/ualberta.ca/onekp/; (Wong et al., 2020) for 13 additional orchid species.

### Library preparation, targeted enrichment and sequencing

The quality of the DNA extractions, including concentration and the distribution of fragment lengths, were checked using a Qubit 3.0 Fluorometer (Life Technologies, Carlsbad, CA, USA) and a TapeStation 42000 system (Agilent Technologies, Santa Clara, CA, USA). Genomic library preparation and enrichment were conducted following the protocols of Johnson et al. (2019). Here, dual-indexed Illumina genomic libraries were prepared for each DNA sample using an insert size of ~350 bp and the Ultra II Library Prep Kit (New England BioLabs, Ipswich, MA, USA), following the manufacturer’s protocol. We used the Angiosperms353 bait-kit to enrich each genomic library in 353 low-copy nuclear genes (Johnson et al., 2019; https://arborbiosci.com/genomics/targeted-sequencing/mybaits/mybaits-expert/mybaits-expert-angiosperms-353/). Sixty-two genomic libraries were pooled in equimolar quantities, to make a 1ug (total DNA) hybridisation reaction, which was subsequently cleaned and sequenced on an Illumina MiSeq v3 (600 cycles, 300 bp paired-end reads) at the Royal Botanic Gardens, Kew to produce ca. 14.8 Gigabase pairs (~30 million paired-end reads).

### Plastome phylogenomics

To assess the performance of the Angiosperms353 nuclear loci for resolving orchid relationships and investigate nuclear/plastid gene tree discordance at different taxonomic levels, we utilized the 78-plastid-gene dataset of (Serna-sánchez et al., 2020): 264 species representing 117 genera, 28 subtribes and 18 tribes. The taxon sampling of nuclear and plastid datasets overlaps in 34 genera, 14 tribes and 22 subtribes. Detailed information on the completeness of this plastid DNA dataset is provided in Serna et al. (2020).

### DNA sequence data analysis and phylogenomic inference

The quality of the newly generated sequencing data was assessed using the FastQC software (freely available at https://www.bioinformatics.babraham.ac.uk/projects/fastqc/). Paired-end DNA sequencing reads were adapter-trimmed and quality filtered with the pipeline TrimGalore! v.0.6.5 (freely available at https://www.bioinformatics.babraham.ac.uk/projects/trim_galore/) using a Phred score quality threshold of 30 (flag -*q*), a minimum read length value of 20 (flag --*length*) and retaining only read pairs that passed all quality filtering thresholds. Data obtained from the SRA were already adapter- and quality-filtered. For each sample, the Angiosperms353 loci were retrieved through the pipeline HybPiper v.1.3.1 (Johnson et al., 2016) by mapping the clean reads against template sequences of the 353 low copy nuclear genes (available at https://github.com/mossmatters/Angiosperms353) using the Burrows-Wheeler alignment (BWA) program v.0.7 (Li and Durbin, 2009) and then *de-novo* assembling mapped reads for each gene separately using the software SPAdes v. 3.13 (Bankevich et al., 2012), with a minimum coverage threshold of 8x. For each gene, homologous sequences from each species were combined and aligned with the software MAFFT v. 7.4 (Katoh and Standley, 2013) using the FFT-NS-i strategy.

For each nuclear or plastid gene alignment, a Maximum Likelihood (ML) gene tree was computed using the software RAxML v8.0 (Stamatakis, 2014) with the GTR+GAMMA nucleotide substitution model, and node support was calculated using 500 bootstrap replicates. The same approach was used to build a nuclear species tree based on a supermatrix made of the 353 concatenated nuclear gene alignments, and to build a plastid species tree based on a supermatrix made of the 78 concatenated plastid gene alignments. To account for topological incongruence between gene trees, we also inferred a nuclear species tree by analysing the ML nuclear gene trees together under the multispecies-coalescent (MSC) framework implemented in the software ASTRAL-III v5.6 (Mirarab and Warnow, 2015). The same approach was used to produce a MSC plastid species tree based on the plastid gene trees. In both cases, branches with Likelihood Bootstrap Support (LBS) > 20 in the gene trees were first collapsed using the software Newick Utilities toolkit (REF), as recommended by (REF). The resulting MSC topologies were annotated with quartet support values for the main topology (*q1*), the first alternative topology (*q2*), and the second alternative topology (*q3*; flag -*t* 2). In addition, we used the software SplitsTree4 (Huson, 1998) to infer Neighbor-net networks based on uncorrected P-distances calculated from the nuclear supermatrix. Neighbor-nets (also known as split-graphs) are suitable diagrams to represent evolutionary relationships in groups that have experienced reticulation (Rutherford et al., 2018) and are useful to identify relationships that exhibit some ambiguity (Solís-Lemus et al., 2017).

### Quantification of intragenomic and nuclear-plastid discordance

The proportion of intragenomic discordance in nuclear and plastid datasets was evaluated by looking at the normalised quartet scores produced by ASTRAL III when inferring MSC plastid and nuclear species trees. The quartet score indicates the proportion of gene tree quartets that are in agreement with the species tree and its magnitude is inversely proportional to incongruence, where a value of 1 indicates potential absence of gene tree discordance.

To assess the degree to which the evolutionary history reflected by the nucleus tracked that of the plastid genome, we compared Euclidean distances among terminals between each bootstrap replicate of each nuclear ML gene tree and each bootstrap replicate of the supermatrix plastid ML tree. The supermatrix plastid ML tree and their bootstrap replicates were chosen over individual plastid ML gene trees because of the high degree of intragenomic congruence observed between the topologies derived from individual plastid loci (see *Results*). The comparisons were conducted with the Procrustean Approach to Cophylogenetics (PACo) pipeline implemented in the R software (Balbuena et al., 2013). The pipeline, originally designed to investigate cophylogenetic patterns between host and parasites, assesses the similarities between any two given trees by comparing the Euclidean distances separating terminals in both via Procrustean superimposition. The efficiency of PACo to assess tree incongruence between nuclear-plastid associations was previously evaluated by (Pérez-Escobar et al., 2016), and the pipeline has been widely used in other plant groups including Compositae (Vargas et al., 2017), Fagaceae (Yang et al., 2018), Orchidaceae (Pérez-Escobar, 2016; Pérez-Escobar et al., 2016) and Rosaceae (Morales-Briones, Romoleroux, et al., 2018). The pipeline provides the sum of squared residuals for each pair of terminals (i.e. association) and each pair of topologies evaluated (Balbuena et al., 2013). This sum of squared residuals can be interpreted as a concordance score because it is directly proportional to the magnitude of the topological conflict for the pair of terminal considered.

Because extremely long branches can bias the comparison of phylogenetic distances between terminals (De Vienne et al., 2011), we conducted PACo analyses on cladograms by assigning a value of 1 to each branch length in each tree, using the function *br.length* of the R-package APE (Paradis et al., 2004). Differences in the position of terminals between nuclear and plastid trees was summarised in barplots using the R-package GGPLOT2 (available at https://ggplot2.tidyverse.org/), for which the sum of square residuals for each pair of terminals across nuclear genes was classified in quartiles. Here, the magnitude of the discordance was assessed by the proportion of genes binned in quartiles 3 and 4 (50% and 75%) in each terminal: the more genes binned in quartiles 3-4, the more discordant the terminals.

To test for intergenomic conflicts occurring across different taxonomic levels, we conducted the same analysis but with trees including one representative of each genus sampled in our nuclear and plastid phylogenomic trees. For this, we conducted sequence alignments and ML inference on each nuclear gene alignment and nuclear and 78-gene matrices (see *DNA sequence data analysis and phylogenomic inference* of *methods*). We then produced subtribe and tribe-level trees by keeping one representative of each clade in our nuclear gene and plastid supermatrix trees following the approach of (Matzke, 2013) as implemented in the R-package BioGeoBEARS (script available at http://phylo.wikidot.com/example-biogeobears-scripts#pruning_a_tree). Here, all tips belonging to the same taxon are pruned except the first species in the list of taxa representing each clade. Terminals identified as potentially conflicting were depicted in tanglegrams using nuclear and plastid ML trees derived from matrices sampled to genus level as implemented in the function *tanglegram* of the R-package DENDEXTEND (Galili, 2015). For subtribe- and tribe-level analyses, we relied on pruned trees originally derived from the nuclear and plastid concatenated supermatrices sampled to genus level.

### Assessment of phylogenetic informativeness and support for nuclear and plastid datasets

We compared performance of the Angiosperms353 low-copy nuclear genes and of the 78 plastid genes for resolving orchid relationships in terms of support and phylogenetic informativeness (PI). We first calculated the proportion of nodes across genus-level nuclear and plastid gene trees that fell into nine discrete LBS categories defined by an interval length of 10 (excluding the interval LBS [81-100]). Secondly, we estimated the PI of each single nuclear and plastid gene alignment with regards to all species relationships recovered in the nuclear and plastid ML species trees, respectively. The trees were made ultrametric by assigning an arbitrary age of 1 to their root and of 0 to their tips (Townsend, 2007), using the software PATHd8 (https://www2.math.su.se/PATHd8/) (Britton et al., 2007). Phylogenetic informativeness of the nuclear and plastid genes was computed in PhyDesign (http://phydesign.townsend.yale.edu/; (López-Giráldez and Townsend, 2011), using the HyPhy algorithm recommended for DNA sequences (Kosakovsky Pond et al., 2005) and the ultrametric ML species trees and nuclear and plastid supermatrices with gene partition information as input.

## RESULTS

### Nuclear phylogenomics of Orchidaceae

Success of the target genes enrichment ranged from 5% (*Neottia nidus-avis*) to 87% (*Dendrobium ellipsophyllum*) gene recovery (**Appendix S2, S3**). The proportion of nuclear genes recovered from SRA and 1KP datamined accessions ranged from 16% (*Phalaenopsis equestris*) to 75% (*Mexipedium xerophyticum* and *Paphiopedilum malipoense*). After excluding samples with less than 15 genes retrieved and gene alignments including less than 20 sequences (i.e. ~80% missing taxon sampling), the final nuclear dataset consisted of 294 genes and 89 species (**Appendix S3**).

Split graphs derived from the nuclear supermatrix revealed clear clustering between members of each orchid subfamily, tribe and subtribe (Fig. 1). However, uncertainty regarding the phylogenetic placement of Neottieae, Nervilieae and Xerorchideae representatives was reflected in an increased number of alternative splits connecting these groups to representatives of other subfamilies. Maximum likelihood inference of the nuclear supermatrix and MSC analyses converged on similar, strongly supported topologies (Fig. 2, **Appendix S4**). However, we found important differences between these analyses regarding the placement of Gastrodieae (represented by the fully mycoheterotrophic *Gastrodia elata*), which was placed as (Neottieae (Gastrodieae (Xerorchideae/other epidendroids))) in the coalescent tree, but as (Neottieae (Xerorchideae (Gastrodieae/other epidendroids))) in the ML results.

**Figure 1.**
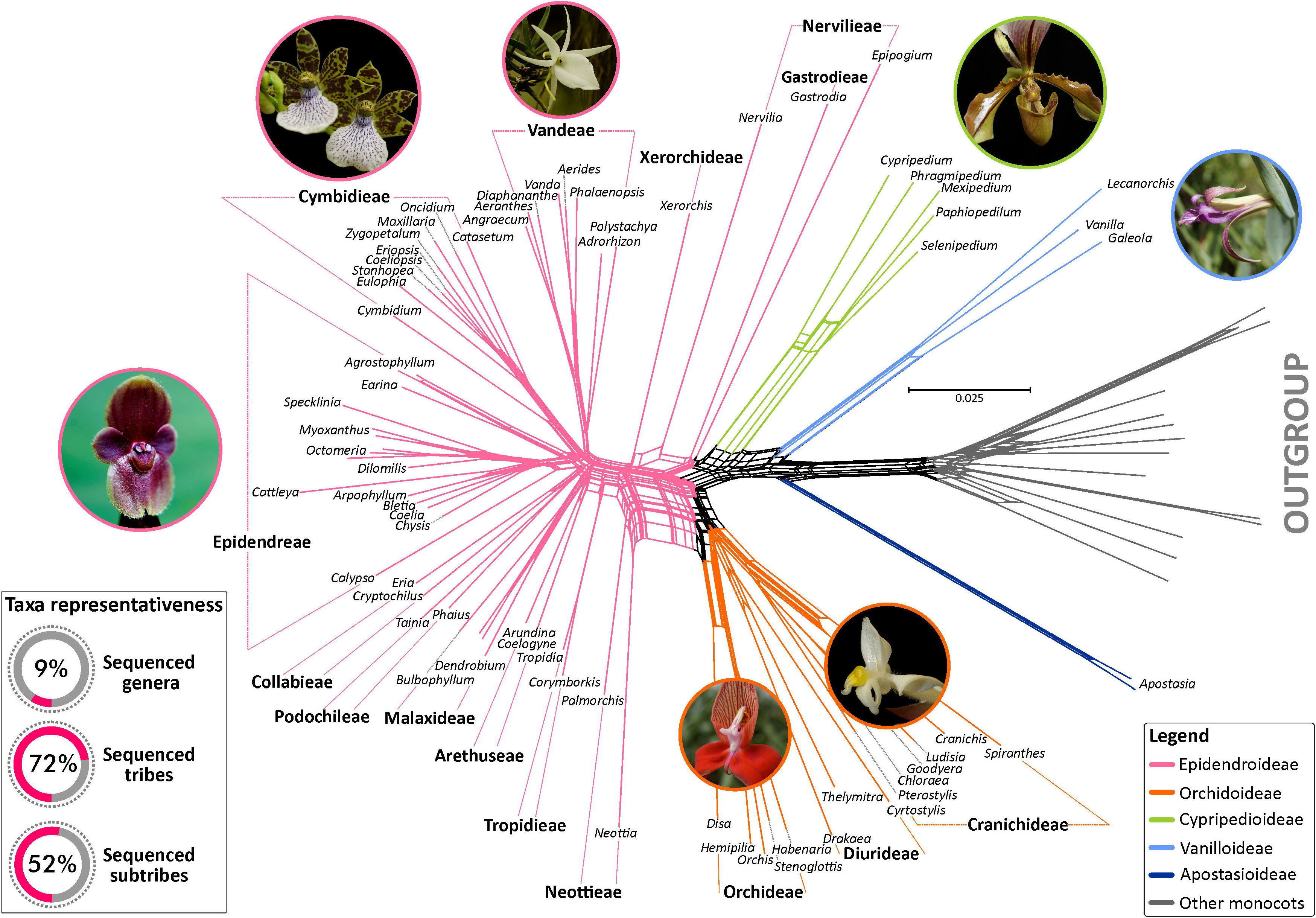
Split network of the Orchidaceae computed from uncorrected P-distances and a concatenated supermatrix of 292 low copy nuclear genes and 75 species. Splits are color-coded by orchid subfamilies (see Legend). Inset box: Taxonomic representation of orchid genera, tribes and subtribes sampled by nuclear gene datasets in this study.

**Figure 2.**
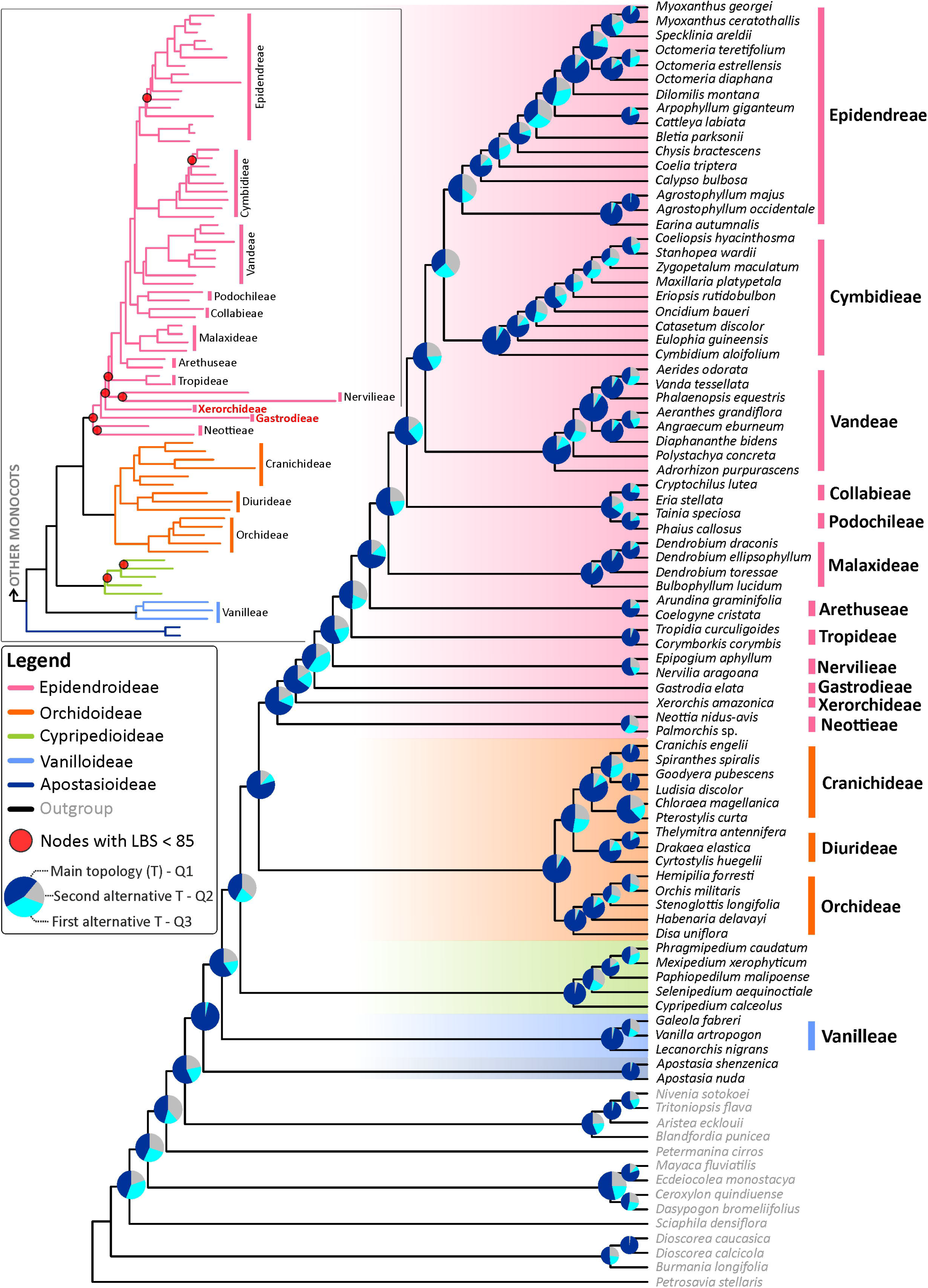
Species-coalescence tree of the orchid family inferred from 292 Maximum Likelihood (ML) gene trees. Pie diagrams at nodes represent quartet support values, with q1 (deep blue portion) representing the proportion of gene tree quartets that support the main (depicted) branch, q2 (blue portion) representing the proportion of quartets supporting the first alternative branch and q3 (grey portion) representing the proportion of quartets supporting the second alternative branch (an explanation of how the quartet values are computed is available at https://github.com/smirarab/ASTRAL/blob/master/astral-tutorial.md). (Inset): ML phylogeny derived from a concatenated supermatrix of 292 low copy nuclear genes. Terminal names with alternative positions to those obtained by the species-tree coalescence analysis are highlighted in bold and red. A detailed version of the phylogeny is presented in Figure S2.

The normalised quartet score of 91% derived from coalescent inference suggests that the majority of gene tree quartets are in agreement with the species tree. Likewise, the quartet support indicated that for most nodes between 40%-96% of the gene trees agreed with the species tree topology. For a few nodes, the proportion of gene trees supporting the quartet displayed in the species tree was below 40%. This included the most recent common ancestors (MRCAs) of *Maxillaria, Zygopetalum, Stanhopea* plus *Coeliopsis* (36.27), *Bletia/* Epidendreae (34.52) and Cymbidieae/Epidendreae (38.76; Fig. 2). In particular, support for the *Neottia/Palmorchis* pair was flagged by ASTRALIII as unreliable due to low numbers of gene trees supporting it (i.e. 12). Likelihood bootstrap support (LBS) values for the ML orchid tree were high, with only 15 branches displaying LBS <100, of which nine had LBS < 85 (**Appendix S4**). The majority of branches with low support were located near the base of Epidendroideae, in line with our findings of nodes with quartet support values < 40 as inferred by the MSC estimation.

### Intragenomic conflict in nuclear and plastid datasets

Estimation of the proportion of gene tree quartets that agree with the species tree through normalised quartet scores indicated that intragenomic incongruence was low. Here, the proportion of gene tree quartets in agreement with the species tree was 89% in nuclear trees and 95% in plastid trees. The proportion of gene quartets supporting topologies other than the species tree (i.e. quartet supports) revealed that the majority of nodes obtained values between 40-95% for the main tree (*q1*) in MSC analyses conducted on nuclear and plastid datasets (**Appendix S5**). Exceptions to this pattern in the nuclear dataset were a few nodes with quartet values supporting alternative bipartitions linked to the MRCAs of Vandeae/Cymbidieae (*q1*=38.6; *q2*=38.6; *q3*=22.69) and *Gastrodia*/remainder of epidendroids (*q1*=40.3; *q2*=45.2; *q3*=14). In the plastid dataset, only four nodes obtained quartet values robustly supporting multiple quartets. Three were in Pleurothallidinae and one represented the MRCA of Podochilieae/Collabieae plus remainder of Epidendroideae (**Appendix S5**).

A similar trend was found in the proportion of gene trees supporting the species tree or alternative topologies in the nuclear dataset sampled to genus level (**Appendix S6**), in which the proportion of gene tree topologies congruent with the MSC species-tree nodes ranged from 15% to 75% and was often dominant over the proportion of gene trees supporting alternative topologies. Three notable exceptions to this pattern were the MRCA of Cymbidieae/Vandeae and two early divergent nodes in Epidendroideae: MRCA of Nervilieae/remainder of Epidendroideae and Malaxideae/remainder of Epidendroideae. Here, gene trees supporting a second most common topology were dominant over the species-tree topology (**Appendix S6**). Overall, the proportion of gene trees supporting the species-tree topology in early divergent nodes in the Epidendroideae was low (from 21-50) but dominant over the proportion of gene trees supporting the second most common topology and any other.

### Nuclear-plastid phylogenetic discordance

The incongruence analysis conducted in PACo on nuclear and plastid trees suggested ten terminals as potentially conflicting (i.e. terminals with ~50% of their squared residual values falling into quartiles 3-4; see *Methods;* Fig. 3, **Appendix S7, S8**). The square residual values of these terminals computed individually for each nuclear gene tree assigned to quartiles 3-4 were overall higher compared with non-conflicting terminals (Fig. 3; **Appendix S8**). After inspecting the position of these terminals in nuclear and plastid ML and MSC trees, *Cattleya*, *Coelia*, *Calypso* and *Earina* appeared to be conflicting with moderate-strong (LBS 85-100) to weak-strong support (LBS 40-98) in the nuclear and plastid ML trees, respectively (**Appendix S8, S9**). These terminals were further linked to nodes with quartet 1 (*q1*) values > 40 < 76 in the coalescent tree inferred from the nuclear dataset (Fig. 3, **Appendix S5, S8**). In the plastid dataset however, the coalescent tree did not recover the same topology as did ML, placing *Earina* and *Coelia* as the sister terminals to Epidendreae with *q1>* 50 (**Appendix S5**), *Calypso* as sister to *Changnienia (q1=* 60), and *Cattleya* as sister to Pleurothallidinae (*q1*=73). Five other terminals identified as potentially conflicting (i.e. *Angraecum, Catasetum, Eulophia, Phalaenopsis and Vanda*) were found in discordant positions with strong support in nuclear and plastid ML and coalescent trees (LBS > 88; *q1* > 48 < 76; Fig. 3, **Appendix S5, S8, S9**).

**Figure 3.**
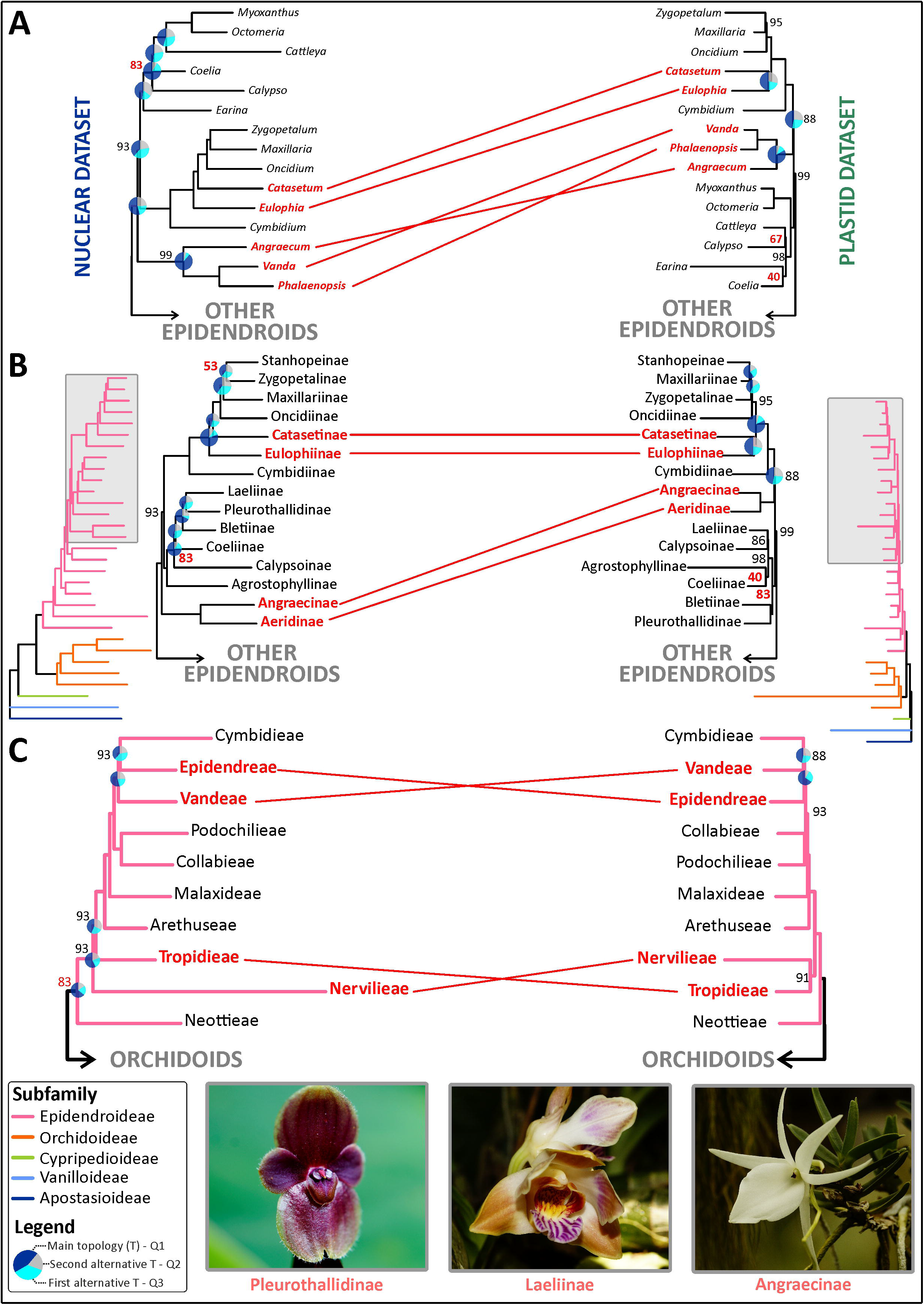
Summary of nuclear-plastid phylogenomic incongruence based on PACo analysis across different taxonomic levels between lineages of the Epidendroideae subfamily depicted based on nuclear and plastid ML trees. Conflicting positions between orchid A) genera, B) subtribes and C) tribes. Association of terminals found to be incongruent and placed with robust support in nuclear and plastid phylogenies are highlighted in bold and red. Pie diagramas at nodes represent quartet support values. Likelihood Bootstrap Support (LBS) values at nodes are 100 unless shown otherwise (LBS < 85 are highlighted in red). (Inset): Three representatives of groups deemed to be conflicting in the orchid family (Pleurothallidinae: *Pleurothallis perryi*.; Laeliinae: *Chysis bracteata*; Angraecinae: *Angraecum eburneum*). Photos: O. Pérez.

Comparisons of nuclear and plastid trees (subtribe level) revealed 13 terminals were potentially conflicting (**Appendix S10**). As in the genus level analysis, the square residual values of terminals computed for each nuclear tree were overall higher for these taxa than the remainder of the tips (**Appendix S10**). However, only the placements of Catasetinae, Eulophiinae, Angraecinae, Aeridinae, Tropidieae and Nervilieae were found to be conflicting with strong support in both nuclear and plastid ML and coalescent trees (Fig. 3; **Appendix S11**). Lastly, incongruence assessments conducted between trees sampled to tribe level revealed that Cymbidieae, Epidendreae, Nervilieae, Tropidieae and Vandeae were potentially conflicting (**Appendix S12**). The phylogenetic positions of Cymbidieae, Epidendreae, Nervilieae, Tropidieae and Vandeae were found to be conflicting with moderate to strong support (Fig. 3, **Appendix S12, S13**) in nuclear and plastid ML trees. Quartet support for the MRCA of Epidendreae/Cymbidieae in the nuclear dataset revealed almost equal support for three alternative bipartitions (*q1*=38.6; *q2*=38.6; *q3*=22.69; **Appendix S5**).

### Phylogenetic Informativeness and support of plastid and nuclear relationships

Profiles of phylogenetic informativeness (PI) for the nuclear and plastid datasets are shown in **Appendix S14**. Net and per-site PIs were in general higher in nuclear than plastid datasets, with average values of 82 and 0.192 versus 56 and 0.058 for nuclear and plastid datasets, respectively. The highest net PI values in nuclear alignments were attained between 0.2 and 0.6 in an arbitrary time scale *t* of 0 (tips) to 1 (root; see *Methods*), broadly corresponding to the initial diversification of the Vanilloideae, Orchidoideae and Epidendroideae. In contrast, the highest per-site PI occurred at *t*[0.4-0.1], coinciding with the divergence times of most generic clades. Net PI values of plastid datasets overall attained uniform values from root to tips, with a notable decrease between *t*[0.4-0] and *t*[0.2-0] with plastid *ycf1* as an exception to this pattern (**Appendix S14**). Per-site PI of plastid datasets presented a broadly similar distribution pattern to the nuclear PI per-site values, attaining their highest values at time intervals of *t*[0.4-.01].

The distribution of LBP across gene tree branches strongly contrasted between nuclear and plastid datasets (**Appendix S4**). A similar distribution pattern of LBS values across the interval t20 (LBS > 10 ≤ 20) and t80 (LBS > 70 ≤ 80) was observed in nuclear and plastid datasets, with t10 (LBS ≤ 10) and t100 (LBS > 80) scoring the largest values. The number of gene tree branches receiving LBS ≥ 80 (t100) in the nuclear dataset was 1600 (vs. 800 in the plastid dataset), whereas the interval t10 scored 1400 branches (vs. 3100 in the plastid dataset).

## DISCUSSION

### Limitations of plastid-only analyses

With few exceptions (Bateman et al., 2018; Bogarín et al., 2018; Unruh et al., 2018; Brandrud et al., 2020; Pérez-Escobar et al., 2020), previous orchid studies (Cameron et al., 1999; Salazar et al., 2003; Neubig et al., 2012; Givnish et al., 2015; Li, Li, et al., 2019; Serna-sánchez et al., 2020) have relied almost exclusively on plastid datasets. This is because nuclear orchid trees have mostly been inferred from the low-copy *Xdh* (Górniak et al., 2010) and ribosomal ITS (nrITS) region (Freudenstein and Chase, 2015). Historically nuclear genes have been difficult to sequence due to a combination of inefficient amplification (due to degradation of DNA samples and/or large intronic regions), the need to clone amplified products due to paralogy or allelic diversity, and lack of universal PCR primers.

High-throughput sequencing has revolutionised our ability to sequence DNA regions at a genomic scale, and thus enhanced our understanding of phylogenetic relationships by utilising different approaches to gain more sequence data per taxon sampled. One of the most commonly used methods is genome skimming (Straub et al., 2012; Dodsworth, 2015), which focuses on sequencing high-copy genomic partitions because these are present even in low-coverage genomic sequencing. Most notably, this includes the plastid genome, which has been used extensively for phylogenetics. The ease of this method has meant that it has surged in popularity over recent years, in many organisms, but also in angiosperms and orchids specifically (Parks et al., 2009; Edger et al., 2018; Li, Yi, et al., 2019; Kim et al., 2020; Zavala-Páez et al., 2020). This has perpetuated the plastid-marker bias in phylogenetic studies of orchids but also greatly improved the results from these analyses. Nevertheless, it has long been acknowledged that without an assessment of nuclear genes, it would be impossible to evaluate a number of important biological phenomena that have shaped angiosperm evolution and diversification, including hybridisation and polyploidisation, population structure, gene flow and introgression (Rieseberg et al., 1996; Vargas et al., 2017; Schley et al., 2020). Given the reliance on plastid analyses, not just as the basis for classification/taxonomy, but also for studying diversification rates, biogeography and trait evolution, it is important to ask to what degree such results reflect organismal evolution. In fact, choosing nuclear over plastid phylogenetic frameworks to investigate characters across time can radically affect our understanding of the mode and tempo of their evolution. An epidendroid example involves the gains and losses of deceit pollination, a trait thought to be linked with increased speciation and extinction rates in orchids (Givnish et al., 2015): using a plastid tree of the orchid subtribes as a framework, the character appeared to have evolved independently in both Laeliinae and Pleurothallidinae (Givnish et al., 2015). If the same character is optimised on our nuclear tree (Fig. 2), deceit pollination would be inferred to have evolved only once at the MRCA of these two subtribes, but this could be a reflection of the sparse taxonomic sampling of our study.

### Are nuclear and plastid evolutionary histories broadly congruent in orchids?

Our comparative phylogenomic analyses provide, for the first time, a solid evolutionary framework for the orchid family inferred from hundreds of low-copy nuclear genes and a detailed assessment on how much these relationships depart from those previously estimated from plastid DNA. An overview of our current understanding of the phylogenetic relationships of orchids was produced by Chase et al. (2015). The topology provided in this review is highly congruent with studies conducted on entire coding plastid and mitochondrial genomes (Givnish et al., 2015; Li et al., 2019; Serna-Sánchez et al., 2019) as well as with that of low-copy nuclear *Xdh* (Górniak et al., 2010). Overall, our quantitative comparisons of our much larger plastid and nuclear datasets support this view, revealing that there is a high degree of congruence between nuclear and organellar phylogenetic trees in orchids, including the monophyly of the five subfamilies and many of the tribal, subtribal and generic relationships (Fig. 3).

Nevertheless, the topological test for quantification of incongruence conducted here reveals that the positions of a few Epidendroideae groups are potentially conflicting. One particular case is the incongruence of *Coelia* and *Earina*, which fall into moderately to well supported conflicting positions in nuclear ML and MSC analyses (**Appendix S5**). The plastid tree of Givnish et al. (2015) placed these genera as sister (LBS 90), a pattern also recovered in our ML plastid phylogeny albeit with lower support (LBS 40). The MSC analysis in contrast robustly places both genera as successively sister to the rest of Epidendreae, but with *Earina* recovering support for an alternative position supported by 25% of the gene quartets (vs 65% of gene quartets supporting the species tree depicted in **Appendix S5**). *Earina* is often used for calibration in molecular clock analyses, as it is one of the three orchid macrofossils that has been unambiguously identified (Conran et al., 2009), raising the possibility of problems in estimating ages for the orchid tree of life.

Another notable exception to the general plastid-nuclear congruence include the interrelationships of Epidendreae, Cymbidieae and Vandeae. These three clades account for nearly a third of the known orchid species diversity worldwide and are thought to be derived from multiple rapid diversifications (Givnish et al., 2015; Pérez-Escobar, Chomicki, Condamine, Karremans, et al., 2017). Previous plastid phylogenomic studies and our own reconstruction place Epidendreae as sister to Cymbidieae/Vandeae with maximum (Givnish et al., 2015; Li, Li, et al., 2019) to moderate support by (Fig. 3, **Appendix S5, S12, S13**), respectively. In contrast, our nuclear tree places Vandeae as sister to Cymbidieae/Epidendreae with strong LBS (Fig. 3, **Appendix S13**). However, we must be cautious to which extent the entire nuclear genome agrees with such a branching pattern. The quartet support generated for the MRCA of Cymbidieae and Epidendreae indicate that an equal proportion of gene quartets support this and an alternative topology (Fig. 3, **Appendix S5**), which could suggest that nuclear gene tree discordance might be responsible for the equally plausible topologies in the Epidendreae(Cymbidieae/Vandeae) situation. Biological phenomena responsible for gene tree incongruence include incomplete lineage sorting (ILS) and gene flow (Rieseberg et al., 1996; van der Niet and Peter Linder, 2008; Schley et al., 2020). Teasing apart the signal that these phenomena leave on a topology is methodologically challenging (Morales-Briones, Liston, et al., 2018; Morales-Briones, Romoleroux, et al., 2018). Given the overall low proportion of genes supporting the main and alternative topologies (**Appendix S5, S6**), we suggest that increasing the number of genes representing this particular node could help determine if there is a dominant branching pattern overall, as well as the relative influence of ILS and reticulation (Nute et al., 2018).

### Potential of conserved low-copy nuclear genes for orchid phylogenetics

Due to the important caveats of plastid-only analyses, phylogenetics based on nuclear genes has started to become increasingly important over the past ten years. Previously it has been difficult to sequence more than a handful of low-copy genes in any plant group (Dodsworth et al., 2018) but genome complexity-reduction methods such as target enrichment have proven to be efficient for sequencing many nuclear loci (Bogarín et al., 2018; Brewer et al., 2019; Dodsworth et al., 2019; Johnson et al., 2019). This approach enables the capture of hundreds of nuclear genes simultaneously, and these loci are highly variable in their levels of conservation, making them useful across all phylogenetic levels.

Our comparative analyses conducted on the per-gene informativeness and support offered by plastid coding datasets and the Angiosperms353 nuclear bait kit (Johnson et al., 2019) strongly points towards the higher performance of low-copy nuclear genes for resolving phylogenetic relationships in both shallow and deep time. These include the recent divergence of epidendroids and clades that experienced rapid radiations such as Epidendreae/Laeliinae and Arethuseae/remainder of the epidendroids. Groups that remain problematic are those located towards the epidendroid MRCA and those that have been historically difficult to place, including Gastrodieae, Neottieae and Nervilieae (Chase et al., 2016), which proved difficult to sequence in our study (gene recovery was poor; **Appendix S2**). Finding a robust result for such clades will require an increase in their taxonomic representation as well as the number of genes included.

## CONCLUSIONS

We present the first comprehensive assessment of the congruence of nuclear and plastid evolutionary histories of the orchid family and provide a generally robust nuclear phylogenomic framework for the family. Comparative analyses of the performance of hundreds of low-copy nuclear genes versus plastid genes reliably demonstrated that the Angiosperms353 genes in general contain more informative loci and resolve more relationships with higher support. We also discovered that, although the plastid genome largely tracks the evolution of the orchid family as reflected by the similarity of its phylogenetic reconstruction to that produced by our analysis of the nuclear genome, there are a few instances of incongruence at varying taxonomic levels that require further study. This is a clear indication that in spite of the overall congruence between nuclear and plastid data, they are not interchangeable, particularly when it comes to the study of character and trait evolution through time. Both trees are providing insights into orchid phylogeny, and the task before us is not to eliminate one as “flawed”, but rather to seek to integrate them to provide the best possible inferences about the evolution of this vast family. Our study also highlights the benefits of nuclear datasets for assessing the influence of hybridisation and incomplete lineage sorting on patterns of diversification. Here, we found that nuclear gene tree discordance is limited, but nonetheless present and linked to key nodes for understanding the diversification of the orchids and its timing. We predict that our study will lead to research addressing the extent of orchid topological discordance both within the nuclear genome as well as between genomic compartments and the phenomena driving this incongruence.

## ACKNOWLEDGEMENTS

This work was funded by grants from the Calleva Foundation and the Sackler Trust to the Plant and Fungal Tree of Life Project (PAFTOL) at the Royal Botanic Gardens, Kew. OAPE is supported by the Sainsbury Orchid Fellowship at the Royal Botanic Gardens Kew and the Swiss Orchid foundation.

## AUTHOR CONTRIBUTIONS

OAPE, DB, WBB and MWC designed research, OAPE, DB, OM and NE collected samples, OAPE, SB, RC, RS, IK and SKM performed all the lab work, OAPE, DB, SB, MFT and JAB performed all analyses, OAPE, SD, GC, SB and MWC wrote the manuscript, with contributions from all co-authors.

## ONLINE SUPPLEMENTARY MATERIAL

**Appendix S1.** Species names, taxonomic and voucher information (including herbarium specimens whenever available) for material used in this study. The number of Angiosperms353 low copy nuclear genes sampled in phylogenetic estimations and missing data per sample are also provided.

**Appendix S2.** Target enrichment success of *de-novo* sequenced accessions. A) Heatmap denoting the proportion the gene sequence length recovered for each sequenced accession.

**Appendix S3.** Number of reads produced for *de-novo* sequenced accessions. The number of reads mapped and proportion of target gene sequenced are also provided.

**Appendix S4.** Maximum likelihood tree derived from a supermatrix of 292 low-copy nuclear genes. Likelihood Bootstrap Support (LB) is 100 unless shown otherwise (LBS <85 in red). A) Number of branches per LBS interval derived from 292 ML nuclear gene trees. B) Number of branches per LBS interval derived from 78 ML plastid gene tree.

**Appendix S5.** Multispecies Coalescent (MSC) trees inferred from 292 ML nuclear gene trees (A) and 78 ML plastid gene trees (B). Pie diagrams at nodes represent quartet support, with q1 (deep blue portion) representing the proportion of gene tree quartets that support the depicted branch, q2 (blue portion) representing the proportion of quartets supporting the first alternative and q3 (grey portion) representing the proportion of quartets supporting the second alternative. Tribes follow the classification of Chase et al. (2015).

**Appendix S6.** MSC trees inferred from 292 ML nuclear gene trees. Pie diagrams at nodes represent quartet support, with q1 (deep blue portion) representing the proportion of gene tree quartets that support the depicted branch, q2 (blue portion) representing the proportion of quartets supporting the first alternative and q3 (grey portion) representing the proportion of quartets supporting the second alternative. Tribes follow the classification of Chase et al. (2015).

**Appendix S7.** Experimental design of nuclear-plastid incongruence analyses. The maximum number of terminals sampled in genus, subtribe and tribe-level analyses of incongruence are provided together with the corresponding number of nuclear-plastid terminal associations analysed and those deemed potentially conflicting.

**Appendix S8.** A) Conflicting phylogenetic positions between nuclear and plastid trees, depicted on ML trees. Tip names and their corresponding connections are taxa flagged as conflicting. Terminals highlighted in red are placed with strong branch support whole those in grey are weakly supported in either the nuclear or plastid tree. Pie diagrams at nodes are provided only for branches with conflicting terminals. They represent quartet support, with q1 (deep blue portion) representing the proportion of gene tree quartets that support the depicted branch, q2 (blue portion) representing the proportion of quartets supporting the first alternative and q3 (grey portion) representing the proportion of quartets supporting the second alternative. All quartet support values are provided in **Appendix S5**. LBS values are 100 unless shown otherwise (LBS <85 in red). B) Proportion of genes with summary of square residual values binned in quartiles 1-4 for each nuclear-plastid association. Terminals with elevated numbers of genes binned in Q3-4 were deemed potentially conflicting (labelled with an asterisk). Terminal names highlighted in bold and red denote those found to be conflicting with strong support in nuclear and plastid tree.

**Appendix S9.** A) ML phylogeny derived from a concatenated supermatrix of 292 low copy nuclear genes and B) 78-coding plastid genes. LBS are 100 unless shown otherwise (LBS < 85 in red). The color of branches connected to internal nodes denote LBSs.

**Appendix S10.** A) Conflicting phylogenetic positions between nuclear and plastid phylogenies sampled to subtribe level, depicted on trimmed ML trees derived from concatenated supermatrices sampled to genus level. Tip names and their corresponding connections are taxa flagged as conflicting. Terminals highlighted in red are placed with strong branch support whole those in grey are weakly supported in either the nuclear or plastid tree. Pie diagrams at nodes are provided only for branches interacting with conflicting terminals. They represent quartet support values, with q1 (deep blue portion) representing the proportion of gene tree quartets that support the main (depicted) branch, q2 (blue portion) representing the proportion of quartets supporting the first alternative branch and q3 (grey portion) representing the proportion of quartets supporting the second alternative branch. All quartet support values are provided in **Appendix S7.** Experimental design of nuclear-plastid incongruence analyses. The maximum number of terminals sampled in genus, subtribe and tribe-level analyses of incongruence are provided together with the corresponding number of nuclear-plastid terminal associations analysed and those deemed potentially conflicting. LBS values are nodes are 100 unless shown otherwise (LBS < 85 are highlighted in red). B) Proportion of genes with summary of square residual values binned in quartiles 1-4 for each nuclear-plastid association. Terminals with elevated numbers of genes binned in Q3-4 were deemed potentially conflicting (labelled with a black asterisk). Terminal names highlighted in bold and red denote tips found to be conflicting with strong support in nuclear and plastid phylogenies.

**Appendix S11.** A) A subtribe-level trimmed ML phylogeny derived from a concatenated supermatrix of 292 low copy nuclear genes and B) 78-coding plastid genes sampled to genus level. Likelihood Bootstrap Support (LBS) values are nodes are 100 unless shown otherwise (LBS < 85 are highlighted in red). The color of branches connected to internal nodes denote LBS values.

**Appendix S12.** A) Conflicting phylogenetic positions between nuclear and plastid phylogenies sampled to tribe level, depicted on trimmed ML trees derived from concatenated supermatrices sampled to genus level. Tip names and their corresponding connections are taxa flagged as conflicting. Terminals highlighted in red are placed with strong branch support whole those in grey are weakly supported in either the nuclear or plastid tree. Pie diagrams at nodes are provided only for branches interacting with conflicting terminals. They represent quartet support values, with q1 (deep blue portion) representing the proportion of gene tree quartets that support the main (depicted) branch, q2 (blue portion) representing the proportion of quartets supporting the first alternative branch and q3 (grey portion) representing the proportion of quartets supporting the second alternative branch. All quartet support values are provided in **Appendix S7**. LBS values are nodes are 100 unless shown otherwise (LBS < 85 are highlighted in red). B) Proportion of genes with summary of square residual values binned in quartiles 1-4 for each nuclear-plastid association. Terminals with elevated numbers of genes binned in Q3-4 were deemed potentially conflicting (labelled with a black asterisk). Terminal names highlighted in bold and red denote tips found to be conflicting with strong support in nuclear and plastid phylogenies.

**Appendix S13.** A) A tribe-level trimmed ML phylogeny derived from a concatenated supermatrix of 292 low copy nuclear genes and B) 78-coding plastid genes sampled to genus level. Likelihood Bootstrap Support (LBS) values are nodes are 100 unless shown otherwise (LBS < 85 are highlighted in red). The color of branches connected to internal nodes denote LBS values.

**Appendix S14.** Net and per-site phylogenetic informativeness analyses inferred from concatenated A) 292-low-copy nuclear and B) 78-coding plastid gene supermatrices sampled to genus level.

## Notes

### Competing Interest Statement

The authors have declared no competing interest.

